# Evaluation of small non-coding RNAs as a possible epigenetic mechanism mediating the transition from biotrophy to necrotrophy in the life cycle of *Phytophthora infestans*

**DOI:** 10.1101/2021.10.30.466584

**Authors:** Juliana González-Tobón, Alejandra Rodríguez-Jaramillo, Laura Milena Forero, Laura Natalia González, Giovanna Danies, Silvia Restrepo

## Abstract

*Phytophthora infestans*, causal agent of late blight disease of potatoes, causes billion-dollar losses worldwide each year. This plant pathogen is a hemibiotroph, first feeding on the host and later killing it. Even though the transcription dynamics of this transition are characterized, the role that small non-coding RNAs (sRNAs) might have is still unknown. Furthermore, a bioinformatic pipeline to search and analyze sRNAs in *P. infestans*, is needed. Using our proposed pipeline, 146 sRNAs were found to be significantly differentially expressed between the evaluated stages of the pathogen’s life cycle. One hundred of these sRNAs were successfully annotated and classified into nine functional categories. The expression of the genes associated to ten of these sRNAs was validated via qRT-PCR. Among these, the expression levels of genes encoding for effectors were inversely correlated to that of the sRNAs aligning to them, which is expected if sRNAs are indeed regulating their expression. This correlation was not clear for sRNAs in other functional categories and should not be confused with strict causality. This study works as a starting point for considering sRNAs as role players in the transition from biotrophy to necrotrophy in *P. infestans* when infecting *Solanum tuberosum*.

---

*Phytophthora infestans* is a devastating plant pathogenic oomycete that causes late blight disease of tomatoes and potatoes (Schoina & Govers 2015). This pathogen is a hemibiotroph, first feeding on living host tissue (biotrophy) and then continuing to live in dead tissue (necrotrophy) (Fry 2008). It is unclear how and why this transition occurs, but it seems to be related with the secretion of effector proteins by the pathogen to the host, which in turn responds using resistance genes (R genes). Some effectors are expressed early in the infection cycle and others later, obeying the “accelerator and brake” model, which states that they can either slow down (brake) or accelerate necrosis (Zuluaga et al. 2016; Lee & Rose 2010). This creates a coevolutionary arms race between the pathogen and the host.

Gaining or losing an effector can happen in a stable way (mutations) or through flexible mechanisms (epigenetic regulation) (Gijzen et al. 2014; Kasuga & Gijzen 2013). This latter option regulates the expression of a transcript without altering the gene’s sequence, enabling the gene to function in the future. Moreover, this is a fast response that gives the pathogen a plethora of opportunities to respond to its environment. The most studied epigenetic mechanism of *P. infestans* are small non-coding RNAs (sRNAs) (Kasuga & Gijzen 2013; Moazed 2009). These are taught to be involved in transposon control and their length usually ranges between 19-40 bp (varying among species) (Jia et al. 2017; Vetukuri et al. 2012; Fahlgren et al. 2013). *Phytophthora infestans’* genome is repetitive in 75% of its length and sRNAs have been found immersed in these regions. Most interestingly, they usually overlap effector genes (Jia et al. 2017; Vetukuri et al. 2012; Haas et al. 2009). Therefore, it has been hypothesized that they could silence/activate the effectors’ expression by altering how heterochromatized (condensed) these regions are (Gijzen et al. 2014; Moazed 2009; Vetukuri et al. 2012).

The transition from biotrophy to necrotrophy has not been studied much under the light of epigenetics. Available literature on sRNAs focus on their specific types and evolutionary traces in the genome (Bollmann et al. 2016). This may be due to the lack of clear bioinformatic pipelines to analyze this type of data in *P. infestans*. Available pipelines are usually designed for animal or plant data (Rueda et al. 2015; Giurato et al. 2013; Gupta et al. 2012). The aim of this study was to evaluate whether sRNAs can constitute an epigenetic regulatory mechanism for the transition from biotrophy to necrotrophy in *P. infestans*. Additionally, this phenomenon was used as a model to design a bioinformatic pipeline that can be used for the analysis of sRNA data sets in this plant pathogen.

The pipeline used is depicted in Figure 1. Our first step was to *acquire* data (grey bar, Fig. 1). For this, we searched extensively for small RNA-Seq data exclusively from *P. infestans*, whose sequencing libraries did not remove sRNAs, and that included samples of different stages in the pathogen’s life cycle. Only one data set fulfilled these requirements: Asman and collaborators (2014) (Åsman et al. 2014) (GEO accession: GSE63292). The authors briefly inoculated leaves of *Solanum tuberosum* with *P. infestans* and extracted sRNAs at 24, 48, and 72 hours post inoculation (hpi), as well as from mycelia growing on culture media. For their isolate (88069), 24 hpi represented the biotrophic stage, 48 hpi the transition, and 72 hpi the necrotrophic stage (Åsman et al. 2014).

**Figure 1.**
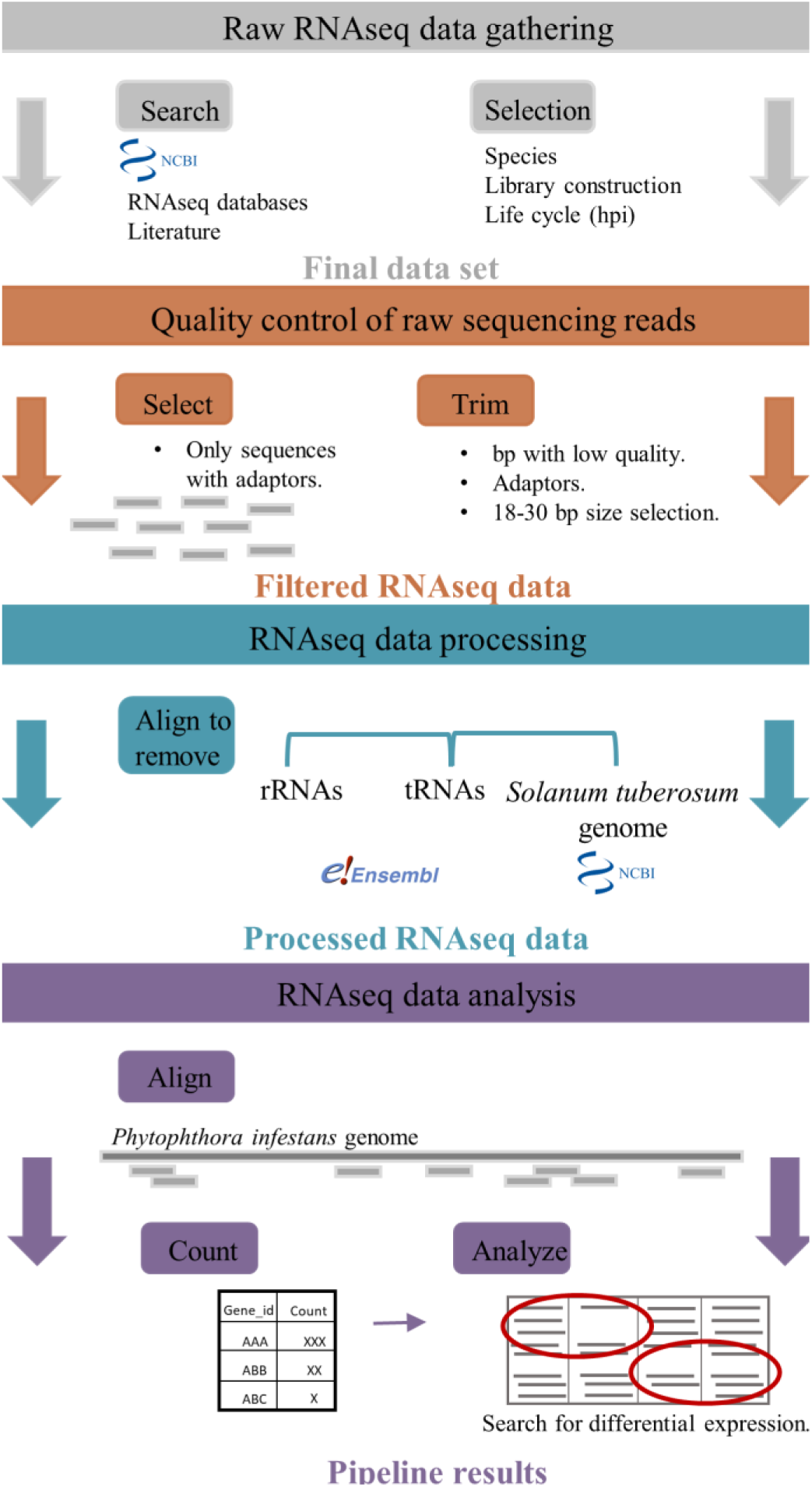
Proposed methodological pipeline for the detection of small non-coding RNAs and analyses of small RNA-seq data. The pipeline follows a sequential order and each colored bar represents one of the four sections of this process. Figure was adapted from Maze et al. 2014.

To *process* this raw data, we performed two big steps: trimming and selection (represented by orange and blue bars in Fig. 1, respectively). We first used Fastq-dump and FastQC to download the data from NCBI and convert it from SRA files to FASTQ format. The reads exhibited very good quality scores (> 40 Phred score) in all base pairs except for the last one, which was removed using cutadapt (Martin 2011). This process was performed only on reads that contained adaptor sequences, which were also removed. Then, only those reads between 18-30 bp were selected using cutadapt to eliminate sequences that were not adequately processed (Vetukuri et al. 2012; Fahlgren et al. 2013; Åsman et al. 2014). All rRNAs (ribosomal RNAs) and tRNAs (transfer RNAs) (Vetukuri et al. 2012) were removed to limit the analyses only to regulatory RNA species. Known rRNA and tRNA sequences of *P. infestans* were downloaded from Ensembl Protists and reads were aligned using Bowtie1 (Langmead et al. 2009). The parameters for these alignments were: only report the first valid alignment (-k 1) and allow 0 mismatches. The resulting reads were aligned to the *S. tuberosum* genome in the same way.

Finally, the reads were ready to be *analyzed* (represented by the purple bar in Fig. 1). They were aligned to the T30-4 *P. infestans* genome from the Broad Institute using Bowtie 1 with the same parameters mentioned above (Vetukuri et al. 2012; Fahlgren et al. 2013; Åsman et al. 2014; Åsman et al. 2016). A GTF file with the annotation of *P. infestans* genes was obtained from Ensembl Protists. This file, as well as the alignment files, were used as input for HTSeq (Anders et al. 2015) to generate count tables that indicated how many times each sRNA was aligned to the genome. These tables were the input for DESeq2 (Love et al. 2014) to determine differentially expressed sRNAs between samples. DESeq2 normalized these counts via the Relative Log Expression (RLE) method (Reddy 2015). This analysis was performed in Galaxy online (Afgan et al. 2016), where the factor specified was “hpi” with four levels: 24, 48, 72, and mycelia in culture. The threshold for p-values associated with the log_2_ fold change (FC) of each sRNA was fixed at 0.05 and each significantly differentially expressed sRNA was annotated using Blast2GO and classified into a functional category.

A set of 146 sRNAs was found to be significantly differentially expressed among the four samples. Log_2_ fold changes for upregulated sRNAs ranged between 2.8 – 4.4, and for downregulated sRNAs between -2 and -4.25. Out of these, six were identified as tRNAs and pseudo tRNAs, and one as a 28S rRNA. Additionally, thirty-nine ncRNAs were associated with putative uncharacterized proteins. The remaining 100 sRNAs were successfully annotated to specific genes and then classified into nine functional categories that are summarized in Figure 2. The first three categories include sRNAs associated with non-protein-coding genes and the remaining ones contain sRNAs mapping to protein-coding genes. Three of the latter included effectors, which are known to be directly associated with the transition from biotrophy to necrotrophy. Therefore, we mostly focused on these three categories for further analyses. A principal component analysis (PCA) among the four individual data sets is shown in Figure 3. As expected, mycelia grown in media differed considerably from any of the other data sets, more than did any of the other data sets between each other. Interestingly, data from 48 hpi and 72 hpi were much more similar between them than when compared to the 24 hpi data set.

**Figure 2.**
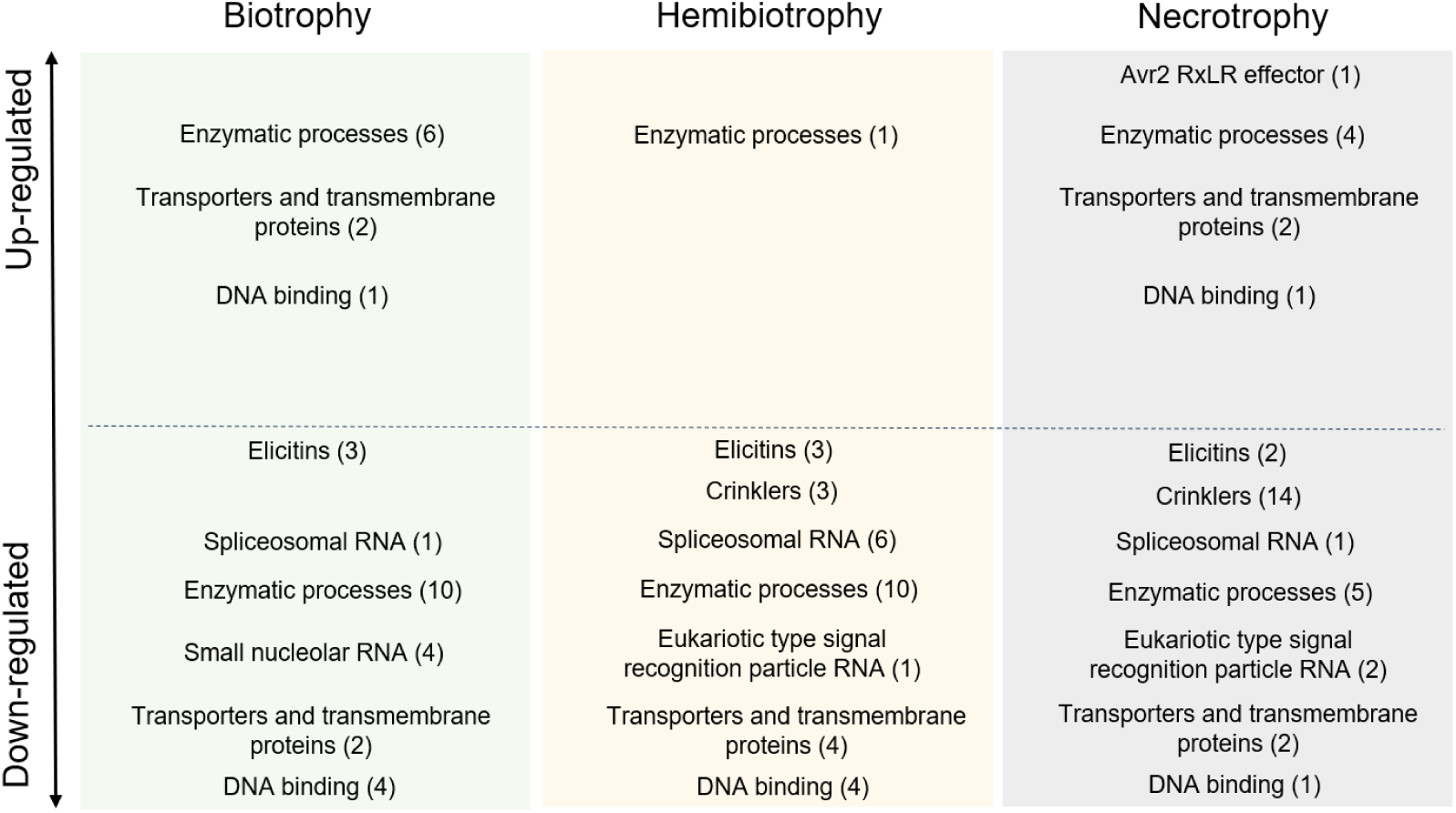
Significantly differentially expressed small non-coding RNAs in *Phytophthora infestans* infecting *Solanum tuberosum* leaves were classified into nine functional categories. Up or down regulated sRNAs belonging to a specific functional category at a specific time point during the cycle are shown. Each box represents one stage of the infection: biotrophy (green), the transition from biotrophy to necrotrophy (yellow) and necrotrophy (grey). Baseline gene expression is calculated in the *in vitro* mycelia, which is represented by the dotted line. Above this line were upregulated, while those below the line were downregulated when compared to the *in vitro* mycelia. The number in parentheses indicates how many sRNAs of each category were either up-regulated or down-regulated at each time point.

**Figure 3.**
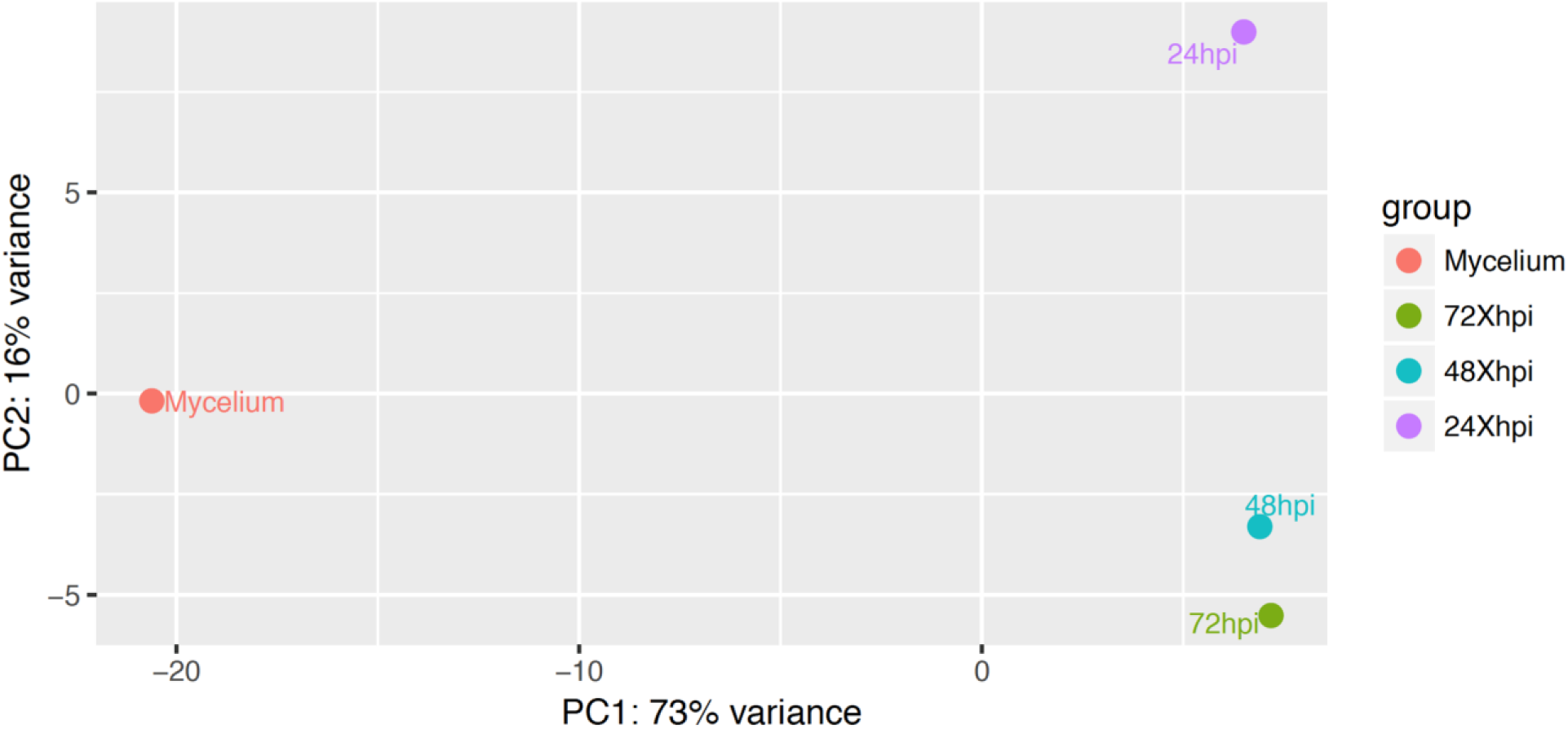
Principal components analysis (PCA) of significantly differentially expressed small non-coding RNAs at each point in the infection cycle. Each dot represents sRNAs of one of the four analyzed data sets (24, 48, and 72 hours post inoculation and mycelia grown in culture media). The distance between them in the grid represent sample-to-sample distances. In other words, how similar these data sets were between them.

To gather further experimental evidence on our predicted sRNAs actually regulating the expression of their corresponding genes during the transition from biotrophy to necrotrophy, the expression of ten of these genes (Table 1) was tested via a quantitative real-time reverse transcription PCR (qRT-PCR). *Solanum tuberosum* plants of the Diacol Capiro variety were grown for one month and a half under greenhouse conditions. Then, they were inoculated with *P. infestans* isolate RC1#10 (EC-1 clonal lineage). The isolate was grown in pea agar for seven days at ± 20 °C. Mycelia was scraped to obtain 1 mL of sporangial suspension and diluted to 1×10^3^ sporangia mL^-1^. Two 10-µL drops of the sporangial suspension were placed on each side of the main vein of the leaf, abaxial side up. Three leaflets per petri dish were maintained in moist chambers until symptoms developed. Subsequently, two transfers to healthy leaflets were performed by immersing the diseased leaves into distilled water and adjusting the solution to 1×10^3^ sporangia mL^-1^ and repeating the inoculation process. This was done to obtain sporangia that came directly from the leaf and not from medium

**Table 1.**
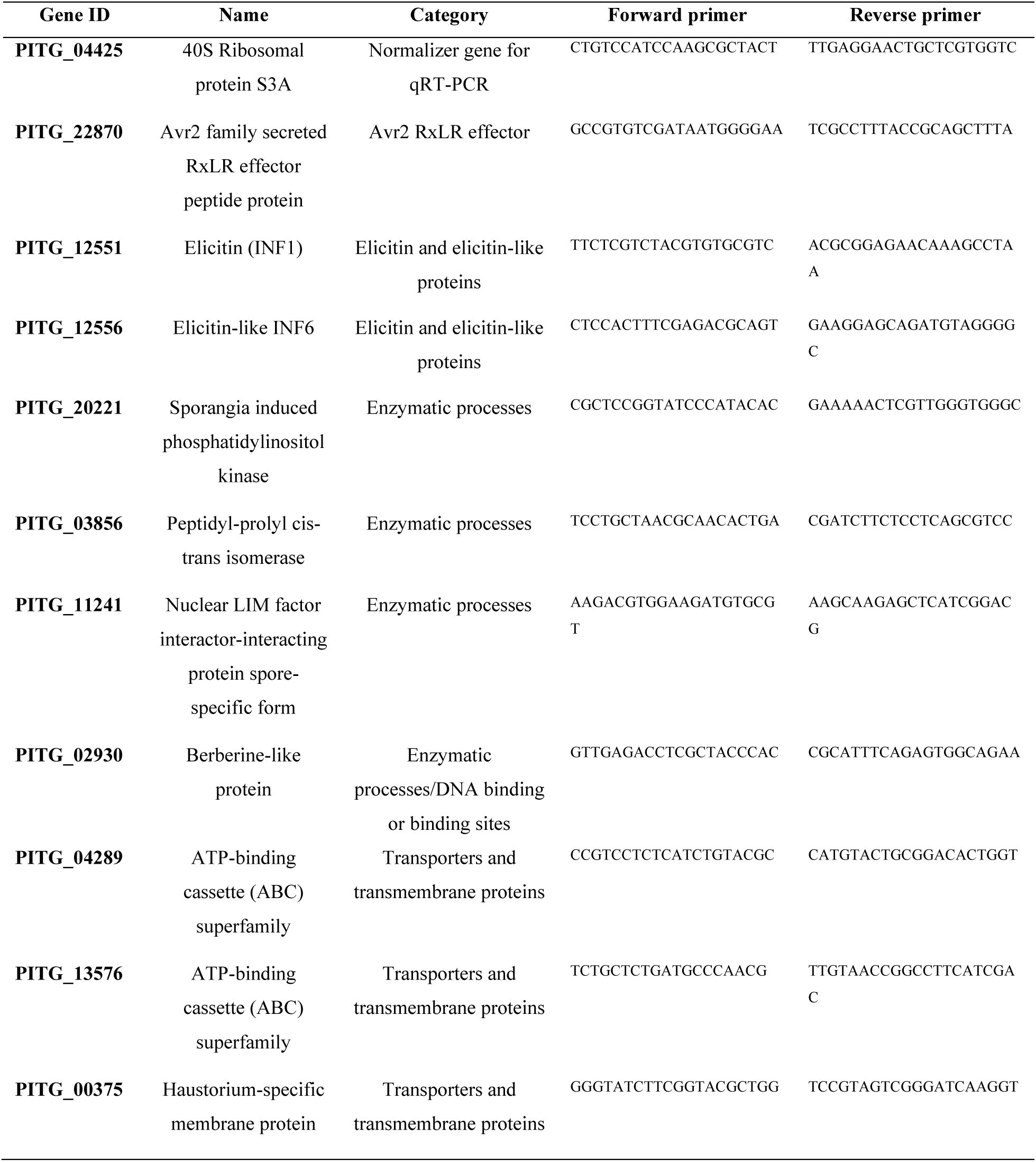
Genes whose expression was tested via qRT-PCR as well as the exon to exon primers designed and used for these assays.

Plant tissue was sampled at 12 hpi (biotrophic stage), 48 hpi (transition from biotrophy to necrotrophy), and 96 hpi (necrotrophic stage). These time points were determined specifically for this isolate and potato variety, under the assessed conditions. The area where each droplet was placed was cut out and frozen at -80 °C. Sporangia from mycelia growing in pea agar were collected as a control and stored at -80 °C. Total RNA was extracted from these using the RNeasy Plant Mini Kit (QIAGEN, Hilden, Germany) following the manufacturer’s instructions. RNA quality was assessed in an agarose gel and quantity was determined using a NanoDrop 1000 Spectrophotometer (Thermo Fisher Scientific, Waltham, MA, USA). Subsequently, extracted RNA was converted to cDNA using RevertAid H Minus First Strand cDNA Synthesis Kit (ThermoFisher), following the manufacturer’s instructions.

Total transcript levels were determined using the Luna Universal qPCR Master Mix (2X) (New England BioLabs, Ipswich, MA, USA), following the manufacturers’ protocol. Gene primers were designed exon to exon using Primer-BLAST software (Ye et al. 2012) (Table 1). The *P. infestans* 40S ribosomal protein S3A (PITG_04425) was used as the constitutively expressed endogenous control as suggested by Yan & Liou (2006). All genes were assayed in triplicate in MicroAmp™ Fast Optical 96-Well Reaction Plates (Applied Biosystems, Foster City, CA, USA), and three biological replicates of each treatment were performed. A control lacking template and a control lacking reverse transcriptase were included. RNA from *P. infestans* grown in culture was used as the calibrator. The 7500 Fast Real-Time PCR System (Applied Biosystems, Foster City, CA, USA) was used and results were analyzed using the REST software (Pfaffl et al. 2002). Results of these qRT-PCR profiles are summarized in Figure 4.

**Figure 4.**
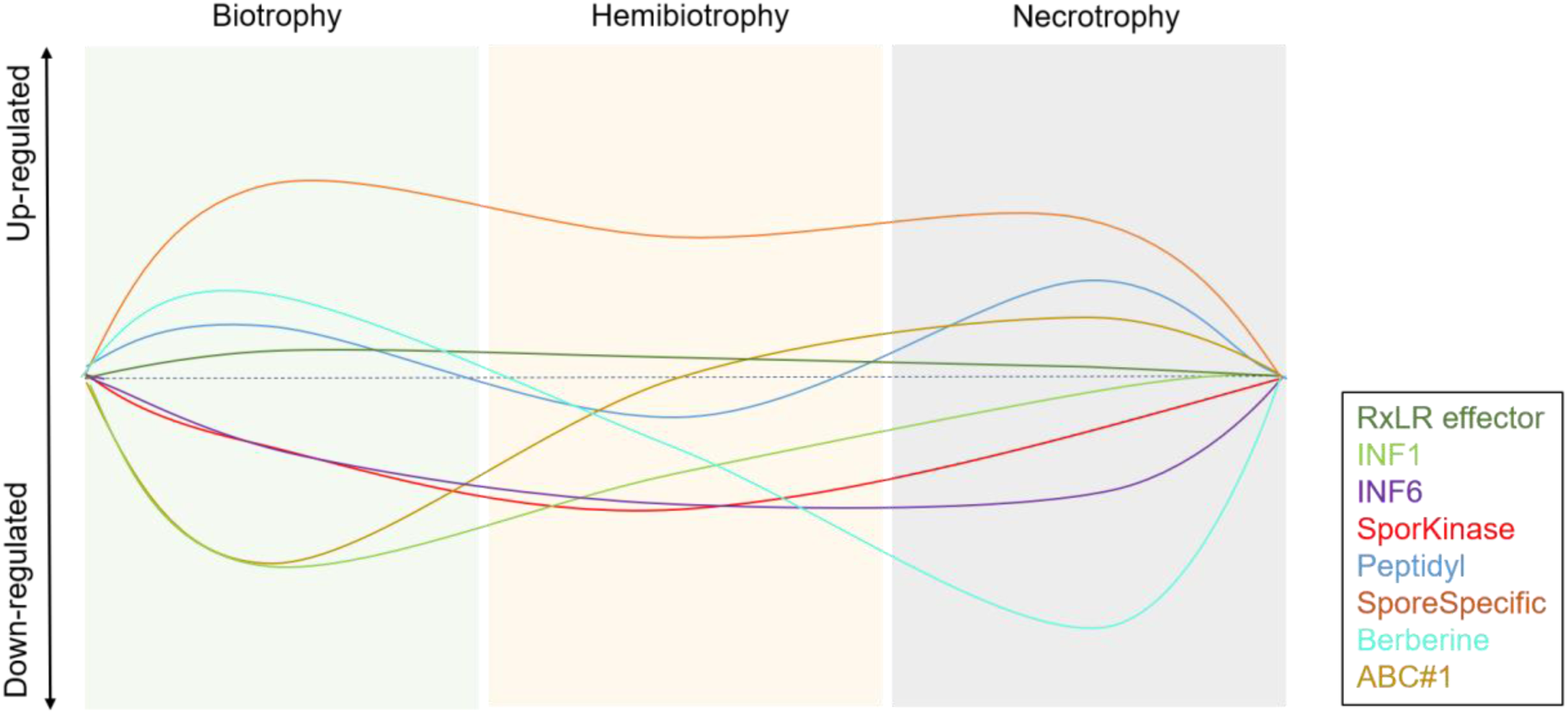
Expression levels identified by qRT-PCR of ten genes associated to relevant small non-coding RNAs during the cycle. Each colored box represents one stage of the infection: biotrophy (green), the transition from biotrophy to necrotrophy (yellow) and necrotrophy (grey). Furthermore, each colored line represents each of the genes that were evaluated (see legend). Baseline gene expression is calculated in the *in vitro* mycelia, which is represented by the dotted line. Genes above this line were upregulated, while those below the line were downregulated when compared to the *in vitro* mycelia.

Three main functional groups were studied in detail: effector-related genes, enzymatic processes and transporter-like proteins. Among the effector genes, one RxLR effector and two elicitins (INF1 & INF6) were tested. The Avr2 family secreted RxLR effector was upregulated during biotrophy and downregulated later in the cycle. Inversely, its sRNA was significantly upregulated at the necrotrophic stage. No other significant differences in its expression were detected during the rest of the cycle (see Figure 5). Since, RxLR effectors are almost always related to the biotrophic stage of the infection this finding was expected (Zuluaga et al. 2016). Furthermore, these results support the proposition that the RxLR sRNA is modulating the gene’s expression by the end of the infection cycle. Similarly, sRNAs associated to crinkler and crinkler-like proteins, usually related to necrotrophy in *P. infestans*, were downregulated at the later stages of infection. This supports the idea that the corresponding genes are generally expressed during these later stages (Zuluaga et al. 2016, Stam et al. 2013). However, the expression of the crinkler genes we used in this study, was not validated using qRT-PCR. Finally, elicitins and elicitin-like proteins are relevant effectors expressed at different time points of the infection. Some are known to be highly expressed during biotrophy while others are highly expressed during necrotrophy (Zuluaga et al. 2016). INF-1, INF-6 and INF-2A are defined as important necrotrophic elicitors (Zuluaga et al. 2016; Kanneganti et al. 2006) and as inducers of various degrees of the hypersensitive response or programmed cell death in plants (Kamoun 2001). However, INF-1 and INF-6 genes were downregulated at all stages of the cycle. The sRNAs associated to them, and to INF-2A, showed the same pattern, thus not supporting the expected inverse correlation (see Figure 5).

**Figure 5.**
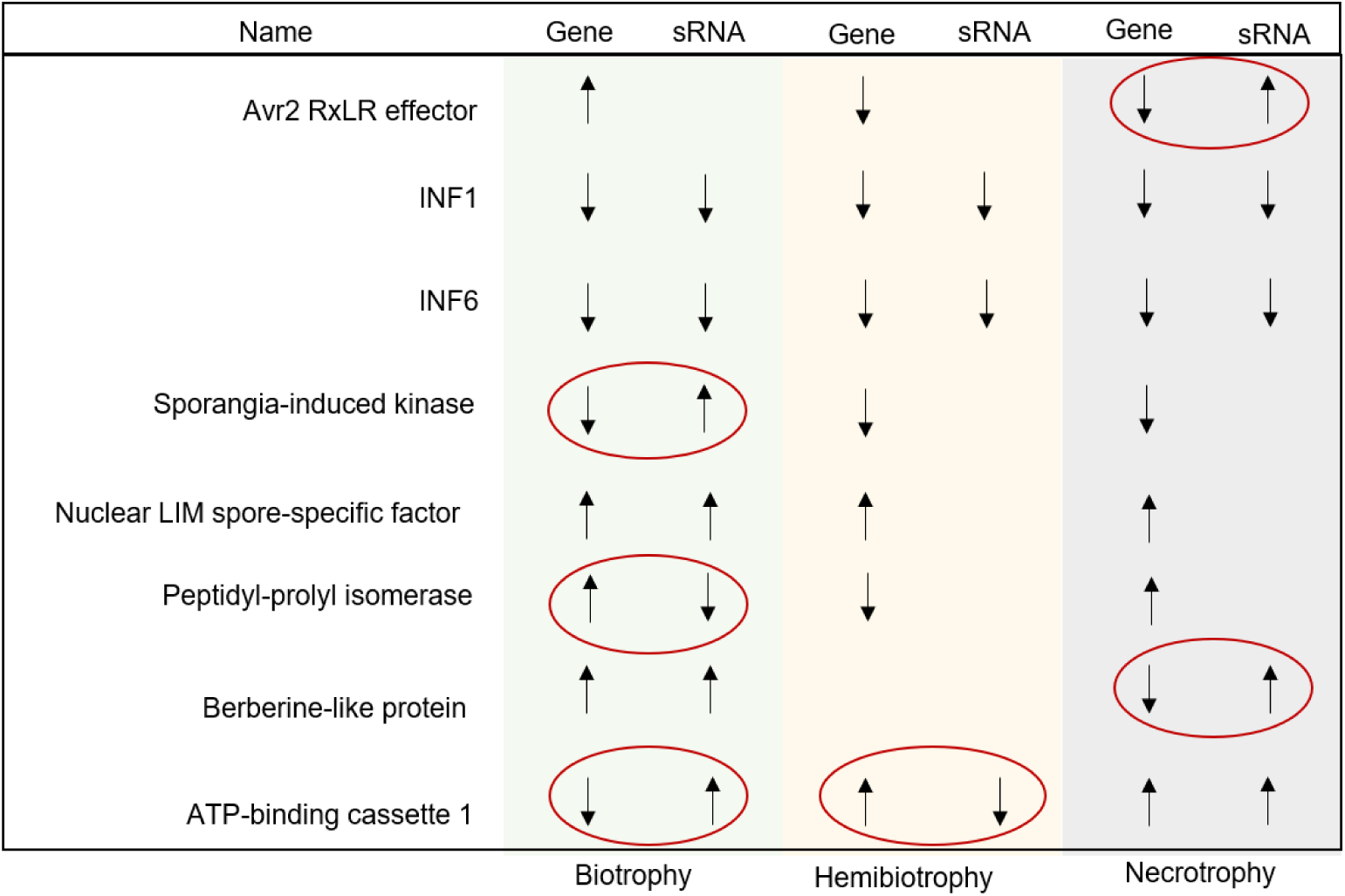
Comparison between significantly differentially expressed genes and their corresponding small non-coding RNAs at different time points during the infection cycle. Eight of the ten genes that were evaluated by qRT-PCR were significantly differential expressed at some point in the infection cycle. Each colored box represents each of the stages in the cycle (green for biotrophy, yellow for the transition to necrotrophy and grey for necrotrophy). Arrows pointing upwards represent upregulation and arrows pointing downwards represent downregulation, for both the gene and its corresponding sRNA. In the cases were no arrow is present for a specific sRNA, no significant differential expression was detected during that specific time point in our bioinformatic analyses. Inverse correlations between the expression of a gene and its sRNA are indicated with a red circle, at relevant time points.

Expression patterns of sRNAs and genes related to enzymatic processes support the regulatory role of sRNAs in most cases (see Figure 5). Two enzymes associated with sporulation and thus expected to express by the end of the infection cycle were tested: a sporangia-induced phosphatidyl inositol kinase and a nuclear LIM spore-specific factor (Tani & Judelson 2006, Judelson & Tani 2007). Only for the kinase gene an expected inverse pattern of expression was detected when compared to its corresponding sRNA during biotrophy (see Figure 5). There was no significant differential expression of this sRNA at other points of the cycle. This could indicate that the sRNA performs a suppressing role at the beginning of the cycle when sporulation has not yet occurred. An inverse pattern was not evident for the LIM spore-specific factor. Additionally, a gene encoding a peptidyl-prolyl cis-trans isomerase also presented an inverse profile when compared with its sRNA during biotrophy (see Figure 5). This highlights the complexity of the disease progression and the regulation of genes involved in it, a process where sRNAs seem to be involved only partly on it.

As for transporters across the cell membrane, we also identified an inverse relation between sRNAs’ and genes’ expression when evaluating a berberine-like protein and an ATP-binding cassette (ABC) transporter. Berberine-like proteins are known to be involved in alkaloid biosynthesis and hydrogen peroxide production. Hence, they are important virulence factors expected to help in disease progression (Meijer et al. 2014). In the same sense, ATP-binding cassette (ABC) transporters are involved in detoxification which allow the pathogen to manage host’s defenses mostly in the form of reactive oxygen species (ROS) (Zuluaga et al. 2016). We did observe an inverse pattern of expression between these genes and their corresponding sRNAs (see Figure 5). For the Berberine-like protein this was observed late in the infection cycle (necrotrophy) and for the ABC transporter earlier in the infection cycle (biotrophy and hemibiotrophy). Finally, the genes for a haustorium-specific membrane protein and a second ABC transporter did not present differential expression in the qRT-PCR assays performed. Testing a larger number of ABC transporters as well as of other transporters and enzymes, might be necessary to understand their expression profiles with more precision.

The expression of sRNAs corresponding to effector genes in *P. infestans* seems to be inversely correlated with these genes’ transcription. This would support the hypothesis of regulatory sequences such as sRNAs performing a role during the pathogen’s transition from biotrophy to necrotrophy. This correlation should not be understood as strict causality since the differential expression observed for these sRNAs could be either a cause or a consequence, which must be further investigated. Moreover, it is important to note that the role of these sRNAs as modulators of specific genes related to infection does not seem to be constant across all the infection cycle but rather localized to certain points within it. Furthermore, sRNAs seem to be involved only partly in regulating the expression of genes related to enzymatic and transportation processes during the disease cycle. However, this study works as a starting point for considering sRNAs as role players in the transition from biotrophy to necrotrophy in *P. infestans* when affecting S. *tuberosum*.

## Acknowledgments

We thank Jose A. Vargas for his valuable advice during the development of this work.

## Declarations

### Funding

No funding was received for this study.

### Competing interests

The authors declare that they have no competing interests.

### Ethics approval

Not applicable.

### Consent to participate

Not applicable.

### Consent for publication

Not applicable.

### Availability of data and materials

The data sets generated and/or analyzed in the current study are available in the GitHub repository https://github.com/jgonzalez10/Small-ncRNAs-pipeline

### Code availability

Code is available in the GitHub repository https://github.com/jgonzalez10/Small-ncRNAs-pipeline

### Author contributions

JG-T developed and implemented the proposed pipeline, analyzed the results, performed all the inoculation steps, the qRT-PCR experiments, and wrote the manuscript. AR-J helped perform the qRT-PCR experiments and the inoculation steps. LMF implemented the pipeline and contributed significantly to the bioinformatic analyses. LNG provided general guidance for the development of the study, specifically for the bioinformatic pipeline implementation. GD provided general guidance for the development of the study and made substantial contributions to writing and editing of the manuscript. SR provided continuous and profound guidance on the development of the manuscript, analyzed the results and was a major contributor in writing the manuscript. All authors read and approved the final manuscript.

## Notes

https://github.com/jgonzalez10/Small-ncRNAs-pipeline

https://www.ncbi.nlm.nih.gov/geo/query/acc.cgi?acc=GSE63292

### Competing Interest Statement

The authors have declared no competing interest.

